# Fate (or state) of CA2 neurons in a mineralocorticoid receptor knockout

**DOI:** 10.1101/2024.11.29.626110

**Authors:** Erin P. Harris, Stephanie M. Jones, Georgia M. Alexander, Başak Kandemir, James M. Ward, TianYuan Wang, Stephanie Proaño, Xin Xu, Serena M. Dudek

## Abstract

Hippocampal area CA2 has emerged as a functionally and molecularly distinct part of the hippocampus and is necessary for several types of social behavior, including social aggression. As part of the unique molecular profile of both mouse and human CA2, the mineralocorticoid receptor (MR; *Nr3c2*) appears to play a critical role in controlling CA2 neuron cellular and synaptic properties. To better understand the fate (or state) of the neurons resulting from MR conditional knockout, we used a spatial transcriptomics approach. We found that without MRs, ‘CA2’ neurons acquire a CA1-like molecular phenotype. Additionally, we found that neurons in this area appear to have a cell size and density more like that in CA1. These finding support the idea that MRs control at least CA2’s ‘state’ during development, resulting in a CA1-like ‘fate’.

## Introduction

The receptors for corticosteroids, glucocorticoid and mineralocorticoid receptors (GRs and MRs, respectively), are important for the regulation of stress responses both in neuronal and non-neuronal tissue^1^. Typically, both receptors can be activated by the circulating stress hormones cortisol or corticosterone and MRs by aldosterone; upon ligand binding, they translocate to the nucleus to operate as transcription factors^2^. Within rodent and human brain, MR (*Nr3c2; NR3C2)* expression is highest in hippocampal area CA2 pyramidal neurons^3–5^. We previously demonstrated that conditional knockout of *Nr3c2* in mice causes a pronounced loss of CA2 marker expression and disrupts CA2’s distinct synaptic properties. In addition, CA2-targeted knockout of MR was sufficient to drive some of the behavioral changes reported with MR knockout in excitatory neurons^5, 6^. These neurons in the CA2 region of the MR knockout mice do not die, but what they become remains unknown, and determining this could reveal insights into the causes of a syndromic autism associated with variants in the *NR3C2* gene^7, 8^.

Prior studies using whole hippocampal isolates have uncovered much of the role of MRs in regulating gene expression, but as yet they have been unable to resolve CA2-specific functions due largely to CA2 being a relatively small portion of the total neurons^2, 9–11^. Successful approaches such as single-cell RNA sequencing^12^ are unlikely to be feasible for CA2-targeted analyses in MR knockout animals because of the loss of many known CA2 cell-type specific gene markers. To begin to bridge this gap, we therefore leveraged the advantages of spatial transcriptomic technology to conduct differential gene expression analysis across hippocampal subfields CA1, CA2, CA3, and the dentate gyrus (DG) of mice with MR conditionally deleted from forebrain neurons (Emx1-Cre: MR fl/fl; MR KO; Supplementary data Fig. 1a). In addition, we analyzed cell size and density in the CA subregions. Together, our findings indicate that the area that would have developed into CA2 takes on a more CA1-like molecular and cellular phenotype.

## Results and Discussion

As evident by expression of several ‘CA2 genes’ like *Amigo2*, *Necab2*, and *Pcp4*, the characteristic molecular profile of CA2 neurons is lost in the MR KO mice (Fig. 1a). In addition, genes like *Prkcb* that are low in CA2 of the wild-type (cre-negative; WT) mice, are expressed at levels similar to that in CA1 and CA3 in the MR KO mice (Fig. 1a, b). That a ‘CA1 gene’, Wolfram syndrome-1 *(Wfs1)*, but not a ‘CA3 gene’, iodotyrosine deiodinase *(Iyd)*, appeared to be expressed in the area that should have been CA2 is strongly suggestive of the CA2 neurons acquiring a CA1-like profile. However, the resolution of the Visium spatial transcriptomics platform is not sufficient to resolve single neurons, so we assessed some of these genes’ expression patterns using single molecule RNA fluorescence *in situ* hybridization (smFISH; Fig. 1b, c). In MR KO tissue, we found evidence of *Wfs1* mRNA in neurons adjacent to *Iyd*-expressing neurons where there would normally be separated by about two-hundred microns. In a few cells, we noted expression of both mRNAs closest to the presumed CA3 border, but largely, the expression of *Wfs1* and *Iyd* was mutually exclusive.

**Fig. 1.**
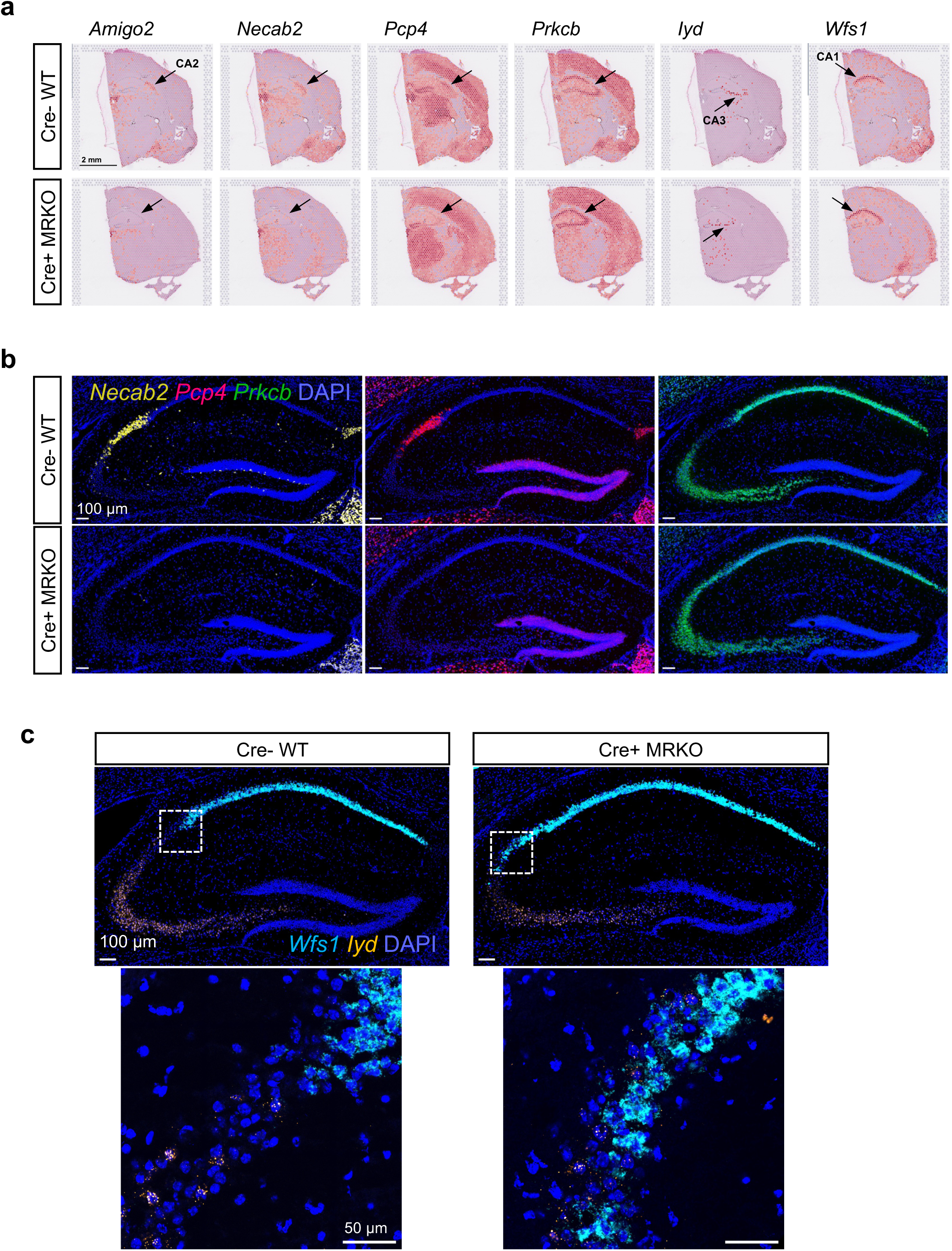
Changes in individual genes with conditional MR KO are evident in spatial transcriptomics and in single neurons. **a**. Examples of brains from a Cre-WT and a Cre+ MR KO mice showing expression of individual genes displayed using the Loupe Browser (10x Genomics). Shown are ‘CA2 genes’ *Amigo2*, *Necab2*, and *Pcp4*, as well as a ‘low in CA2’ gene *Prkcb*. Also shown are the ‘CA1 gene’ and ‘CA3 gene’ *Wfs1* and *Iyd* respectively. Data are not normalized to spot read counts in these displays. Orange/red = higher expression. **b**. Images showing gene expression using single molecule fluorescence *in situ* hybridization (smFiSH) to validate changes in gene expression with MR KO. Shown are *Necab2*, *Pcp4*, and *Prkcb*. Note the loss of *Necab2* and *Pcp4*, and the filling in with *Prkcb* in the area presumed to be CA2 of the MR KO. **c**. smFISH showing that single neurons express neither *Wfs1* nor *Iyd* in CA2 of WT mice. However, neurons in the presumed CA2 region in the MR KO express the CA1 gene *Wfs1*, but not the CA3 gene *Iyd,* with rare examples of neurons expressing both.

These findings again hint at the idea that neurons that would have become CA2 neurons now resemble CA1 neurons, rather than CA3 neurons, in the MR KO mice. However, the ability to make such a conclusion requires the advantages of whole genome transcriptomics, so we sought to assess the entire molecular profile of the area CA2 in WT and MR KO tissue. In this case, we manually selected by location the hippocampal subregions for analysis, as automated clustering based on gene expression profiles would necessarily be altered with MR KO (Fig. 2a, Extended Data Fig. 2). Here we used data obtained from 3 WT male mice (2 cre-negative: *Nr3c2* fl/fl and 1 C57B6J) compared with 3 conditional MR KO male mice (Extended Data Tables 1-3). Similar results were observed with WT and MR KO female mice (1 each was added in Extended Data Fig. 5). Using a principal component analysis (PCA), we found that, consistent with previous data using laser capture microscopy and RNA-seq^13^, the spots in CA2 cluster closer to CA3 than to CA1 in the wild-type tissue (Fig. 2b). However, in the MR KO tissue, the expression profile in spots located in area CA2 no longer cluster with CA3, but instead cluster close to those from CA1 (Fig. 2b). The correlation between CA1 and CA2 (0.810 in Cre-WT) increases in MR KO (to 0.963; Cre+ MR KO) but is unaffected between CA3 and CA2 with MR KO (0.885 in Cre- vs. 0.890 in Cre+).

**Fig. 2.**
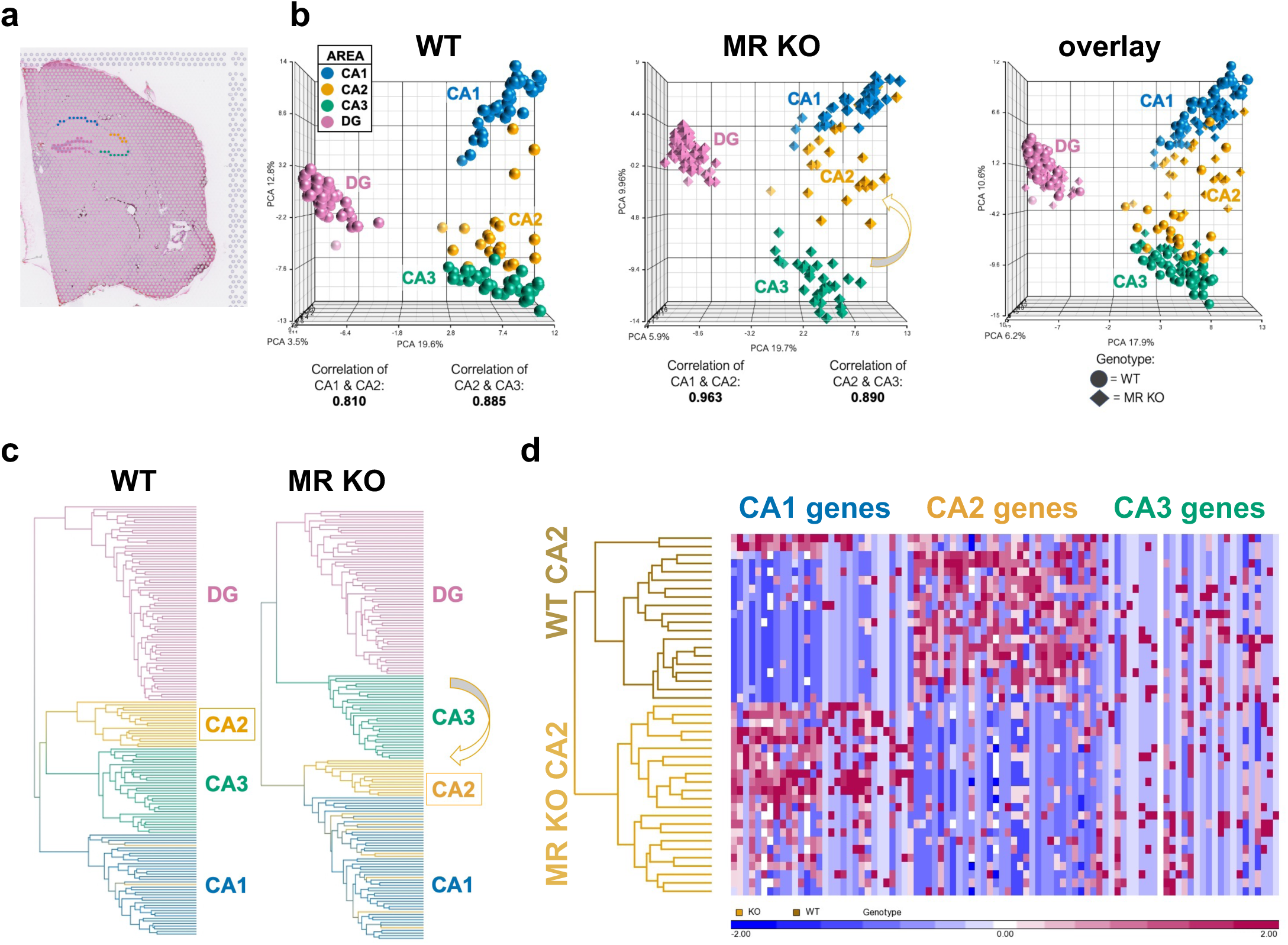
Manual selection of areas allows for analyses of anatomical areas not relying on clustering. **a**. Location of selected spots in CA1 (blue), CA2 (yellow), CA3 (green), and DG (pink). **b**. Principal component analysis (PCA) showing changes occurring with MR KO. Comparison between WT and KO reveals a shift of the transcriptional profile of CA2 in Cre+ MR KO mice towards CA1 as compared with Cre-mice. Correlation between CA1 and CA2 (0.810 in Cre-) increases in MR KO (0.963; Cre+ MR KO) but is unaffected between CA3 and CA2 with MR KO (0.885 in Cre^-^ vs. 0.890 in Cre^+^). **c**. Dendrograms indicating the hierarchical clustering of spots with similar expression profiles based on 368 hippocampal genes (Extended Data Table 1). Lines representing spots are color-coded according to the spot location. **d**. Heatmap of gene expression values for CA2 spots from WT (top) and MR KO (bottom) males for the top 30 CA1 genes, CA2 genes, and CA3 genes as identified by differential expression analysis (Extended Data Tables 2 & 3). Blue = lower expression, Red = higher expression.

We observed similar findings using hierarchical clustering (Fig. 2c, Extended Data Fig. 3a). Spots representing CA2 and CA3, but not CA1, could be clustered by genotype (Fig. 2d, Extended Data Fig. 3b). This analysis also revealed that the changes seen in MR KO were most likely driven by the combined loss of ‘CA2 genes’ and gain of ‘CA1 genes’ with minimal changes to the ‘CA3 genes’ (Fig. 2d). These data strongly suggest that not only are MRs are required for the acquisition and/or maintenance of CA2’s molecular profile, but also that they default to a CA1-like profile in the absence of MRs. Although glucocorticoid receptors (GRs) are upregulated in the MR KO^5^, previous work using a double MR/GR KO still show a loss of CA2 markers^14^, indicating that GRs are not suppressing CA2 gene expression in the MR KOs.

Insights can be gained by looking at the genes up- and down-regulated with MR knockout in the different hippocampal regions. Interestingly, among the CA regions, CA2 had approximately double the number of up- and down-regulated genes than the CA1 and CA3 regions in MR KOs (Extended Data Figs. 4 and 5b). Among the genes significantly reduced in CA2 were the expected ‘CA2 genes’ like *Amigo2*, *Pcp4*, and *Necab2*. Similarly, up-regulated genes in CA2 were some of the ‘CA1 genes’ like *Wfs1* and *Gap43*. The DG however, had by far, the most genes up- and down-regulated in MR KO tissue. In some cases, genes that were highly expressed in both CA2 and DG like *Adcy1* were down-regulated in both regions, suggesting a role for MRs as critical regulators of their expression, whereas some, like *Pcp4*, were unchanged in DG. That said, we found no evidence that the DG was fundamentally altered in its identity distinct from the CA regions. A pathway analysis of genes changed in CA2 of MR knockout tissue revealed networks such as those in NFAT, CREB, GPCR, and synaptogenesis signaling pathways, in addition to circadian rhythm signaling pathways, which might have been expected given the circadian changes in corticosterone (Extended Data Figs. 6a and 7a). Some of the upstream regulators of MR-regulated genes in CA2 showed disease-related genes such as *PRKAA1*, which has been implicated in major depression^15^ (Extended Data Figs. 6b and 7b). Of the significant pathways shared with CA1 and CA3, more in CA1 were shared with those in CA2 (Extended Data Fig. 8).

To investigate further whether MRs are required for the acquisition of structural features of CA2 neurons, we examined cell body density in the different hippocampal areas of WT and MR KO mice. In WT mice, neurons in CA2 and CA3 are larger than neurons in CA1 and so the cell density, as assessed by DAPI stain, is greatest in CA1. Here we measured distance to nearest neighbor and nuclei per µm^3^ (Fig. 3). In both measures, nuclei in CA2 were significantly different from those in CA1, but not different from CA3, in tissue from WT (Cre-negative) animals. This relationship was altered in the MR KO (Cre-positive) tissue in that both distance to nearest neighbor and nuclear density in CA2 were intermediate between CA3 and CA1 (reduced distance between nuclei and increased nuclear density). Both measures were significantly different between WT and MR KO tissue in CA2, but not in CA1 or CA3 (Fig. 3c, d). We interpret these findings to indicate that CA2 neurons, without MRs, come to partially, but not entirely, resemble CA1 neurons.

**Fig. 3.**
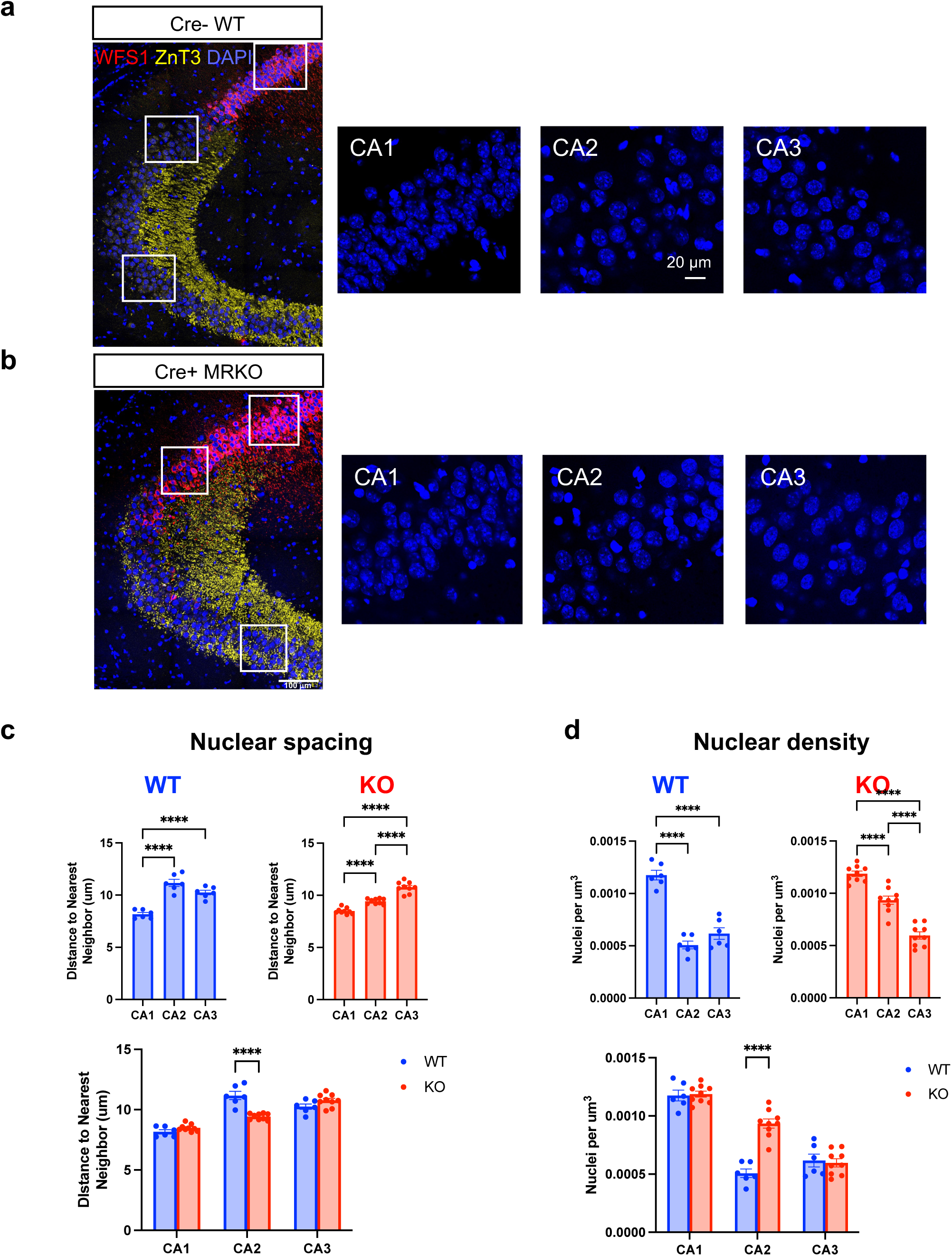
Pyramidal neuron density in area CA2 becomes more CA1-like in MR KO. **a., b**. Immunostaining for ZnT3 to label the mossy fibers and WFS1 to label the CA1 pyramidal cells in WT (a) and MR KO (b) animals were used to localize CA1, CA2 and CA3. DAPI was used to label nuclei. In WT animals, CA2 is positioned at the distal end of the mossy fibers, and WFS1 does not overlap with ZnT3. CA2 was defined as the pyramidal cell nuclei in the pyramidal cell area overlapping with the distal-most 150 um of the ZnT3 stain, and measurements were made from there. In MR KO animals, ZnT3 expression overlaps with the cells expressing WFS1, so CA2 was defined as the area of overlap between these two markers, and measurements corresponding to CA2 were made from that area. Inset squares represent the areas in each subregion that are shown enlarged on the right side. **c**. Nuclear spacing, measured by nearest neighbor analysis, show that in WT animals, both CA2 and CA3 neurons have significantly greater nuclear spacing than CA1, and CA2 spacing does not differ from CA3 (F(2,10)=81.24, p<0.0001, one-way ANOVA; results of Tukey’s post-hoc test shown). In MR KOs animals, both CA2 and CA3 have significantly greater nuclear spacing than CA1, but CA2 has significantly less nuclear spacing than CA3. (F(2,16)=122.3, p<0.0001, one-way ANOVA; results of Tukey’s post-hoc test shown) Nuclear spacing in CA2 was significantly greater in MR KOs than in WTs (main effect of subfield: F(2,26)=164.7, p<0.0001; subfield x genotype interaction: F(2,26)=44.71, p<0.0001; two way-ANOVA, result of Sidak post-hoc test shown). **d**. Nuclear density was measured across the depth of DAPI-stained section. In WT animals, CA2 had a significantly lower nuclear density than CA1 but not CA3 (F(2,10)=97.75, p<0.0001, one-way ANOVA; results of Tukey’s post-hoc test shown, but in MR KO animals CA2 nuclear density was significantly greater than in CA3 (F(2,16)=146.7, p<0.0001, one-way ANOVA; results of Tukey’s post-hoc test shown). Nuclear density was significantly greater in CA2 of MR KO animals than in WT (main effect of subfield: F(2,26)=210.0, p<0.0001; main effect of genotype: F(1,13)=9.863, p<0.001; subfield x genotype interaction: F(2,26)=35.31, p<0.0001; two way-ANOVA, result of Sidak post-hoc test shown), suggesting that CA2 moves toward an anatomical profile more closely resembling CA1 than CA3.

Taken together, these findings are indicative of MR-dependent influences on neuron development, present in CA2, that are not present in CA1 and CA3. These findings have important implications regarding the underpinnings of a syndromic autism caused by variations in the *NR3C2* gene^7, 8^. Further, MRs and CA2 proteins can be down-regulated by corticosterone in rodents^5^, these findings present an avenue by which a part of the hippocampus that regulates social behavior, CA2, could be fundamentally changed during chronic stress.

## Methods

### Spatial Transcriptomics

Brains from 3 EMX Cre-negative and 3 Cre-positive MR KO male mice, aged 3-4 months, were removed following rapid decapitation and frozen in isopentane on dry ice. Tissue was stored at -70° C until use, at which time they were cryosectioned at 10 μm and mounted on Visium slides (10x Genomics, Cat. #1000184). Slides were processed per the 10x Genomics protocol, as used by Vanrobaeys, et al.^17^: tissue was fixed with methanol and stained with H&E followed by imaging. Optimal permeabilization time determined by a tissue optimization kit (10x Genomics, 1000193) was 18 minutes. The on-slide cDNA synthesis and release, cDNA amplification, and library preparation for RNAseq were performed according to the manufacturer’s protocol. Library samples were sequenced on an Illumina NexGen Sequencer (NovaSeq 6000 with read 1 for 28 nt and read 2 for 90 nt length) with an average read depth of 233 million read pairs per brain section, and 86,260 reads and 3,822 genes per spot. Distributions of hippocampal read counts per spot, number of genes per spot, and percent mitochondrial DNA per spot are shown in Extended Data Figure 1b).

For processing of raw spatial transcriptomics data, raw FASTQ files along with slide image for each sample were processed using Space Ranger software (version 1.1, 10× Genomics) to align reads against mouse reference genome mm10. A feature-spot matrix was generated using the Visium spatial barcodes. Seurat (version 4.0) was used to perform clustering analysis of combined dataset^16^. SCTransform was applied to normalize the gene expression values across each spot^17^. Differentially expressed genes (DEGs) were identified by pair-wise comparisons between the 3 WT and 3 MR KO samples within the manually selected spots in CA1, CA2, CA3 pyramidal cell, and DG granule cell layers. Note that although the location of CA2 could be identified in 10xGenomic’s Loupe browser based on gene expression (expression of representative ‘CA2 genes’) and anatomical landmarks, this was not possible for identification of CA2 in the knockout due to loss of such marker gene expression. In this case, anatomical location alone was used to select spots representing CA2. Principal component analysis (PCA) and hierarchical clustering of these spots were generated by the Genomics Suite of Partek software package version 6.6. Volcano plots were constructed using VolcaNoseR^20^. Venn diagrams were generated by Venny 2.1^21^. Differentially expressed genes between Cre-WT and Cre+ MR KO in CA2 with log fold-change values greater than 0.5 or less than -0.5 and FDR <0.05 were included for Ingenuity Pathway Analysis (IPA; Qiagen).

### Single-molecule fluorescent in situ hybridization and imaging

We performed single-molecule fluorescent *in situ* hybridization (smFISH; RNAscope) to validate gene expression changes detected by spatial transcriptomic sequencing and to assess single-cell distribution of gene transcripts. Two animals, one per genotype (Emx1 Cre+/-: MR fl/fl); were sacrificed via rapid decapitation. Brains were extracted and flash-frozen in Tissue Plus® O.C.T Compound (Fisher Scientific, Hampton, NH) in isopentane chilled over dry ice. Brains were stored at -70°C until cryosectioned at 20 μm and mounted on SuperFrost® Plus slides (Fisher Scientific). Tissue sections were probed for target mRNAs according to the manufacturer protocol for the RNAscope Fluorescent Multiplex kit (Advanced Cell Diagnostics, Hayward, CA). Target probes for *in situ* hybridization were Mm-*Nr3c2*-E5E6-C3 (Cat#456321-C3), Mm-*Pcp4*-C2 (Cat#402311-C2), Mm-*Necab2*-C3 (Cat#467381-C3), Mm-*Prkcb*-C1 (Cat#874311), Mm-*Wfs1*-C3 (Cat#500871-C3), and Mm-*Iyd*-C2 (Cat#465011-C2) from Advanced Cell Diagnostics (Hayward, CA). Signal was developed using Opal Dyes 520 (Cat#OP-001001), 570 (Cat#OP-001003), 690 (Cat#OP-001006) from Akoya Biosciences (Marlborough, MA). Sections from both genotypes were processed in parallel and imaged on a Zeiss LSM 880 inverted confocal microscope. Whole hippocampal images were acquired with a Plan-Apochromat 20x/0.8 M27 objective using z-stacks which were collapsed with a maximum intensity projection. 63x images were acquired with a Plan-Apochromat 63x/1.40 Oil DIC M27 objective with a pinhole setting to yield a Z-thickness of 1.7 μm and capture the coexpression of transcripts within the same cells. Acquisition settings were set separately for each staining scheme and held constant across genotypes. Brightness/contrast adjustments were made in FIJI/ImageJ and Powerpoint and applied equally across genotypes for comparison purposes.

### Histology

Six WT and nine MR KO animals were transcardially perfused with 4% paraformaldehyde and brains post-fixed overnight in the same fixative. Brains were sectioned at 40 mm with a vibratome (Leica vt1000) and stored in PBS with sodium azide. For immunohistochemistry, sections were washed in PBS followed by PBS with 0.1% triton X-100 (PBST). Sections were blocked in PBST with 5% normal goat serum for 1 hour followed by overnight incubation in blocking solution plus primary antibodies: rabbit anti-WFS1 (1/500, Proteintech, 11558-1-AP) and guinea pig anti-ZnT3 (1/500, Synaptic Systems, 197 004). Sections were washed in PBST then incubated for 2 hours in blocking solution plus secondary antibodies: goat anti-rabbit 568 (1/500, Invitrogen, A11011) and goat-anti guinea pig 633 (1/500, Invitrogen, A21105). Sections were washed with PBST then PBS, mounted and coverslipped with Vectashield hardset mounting medium with DAPI (Vector Laboratories, H-1500). Sections were imaged on a Zeiss LSM 880 confocal microscope at 40X using tiled Z-scans. Imaris (Oxford Intruments) software was used to perform nearest neighbor analysis and measure total volume of analyzed regions to arise at measures of nuclear spacing and nuclear density. Statistical analyses were performed using GraphPad Prism software.

## Data Availability

The datasets generated and/or analyzed in the current study are available in the Gene Expression Omnibus (GEO), accession number: GSE272919. Original images are available from the corresponding author upon reasonable request.

## Author contributions

SD, GA, and SP conceived of the study. Experiments were performed by EH, SJ, GA, SP, and XX. Data were analyzed by EH, GA, TYW, JW, and BK. SD, EH, SJ and GM wrote the paper with input from the other authors.

## Acknowledgements

We would like to thank Eli Ney of the NIEHS Comparative & Molecular Pathogenesis Branch for scanning the Visium slides and Victor Catalan Gallegos and Priyanka Singh for cutting the brain tissue. This research was funded by the Intramural Research Program of the NIEHS, NIH, ES100221.

## Extended Data Figure Legends

**Extended Data Fig. 1.**
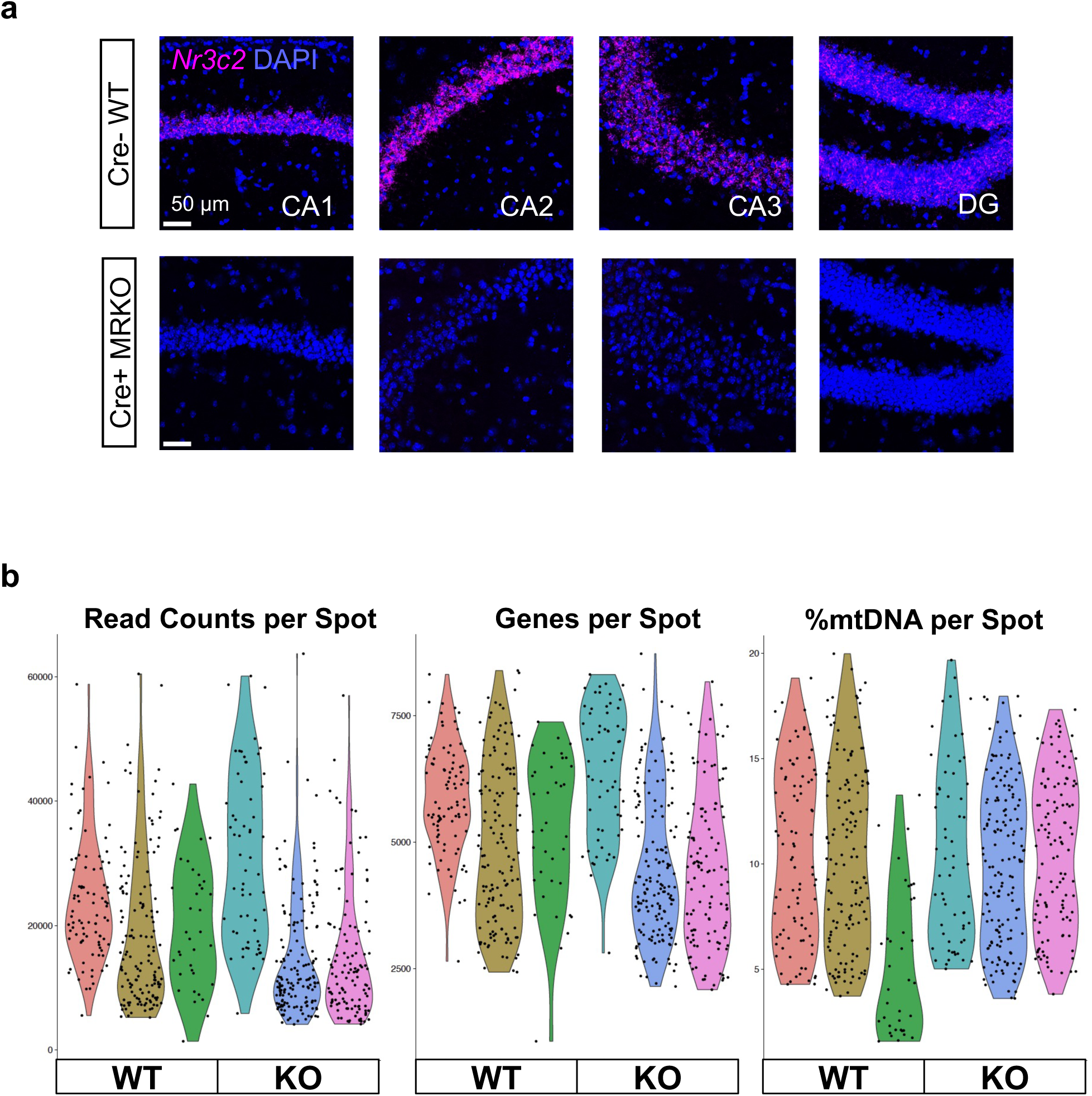
**a**. smFISH for MR (*Nr3c2*) to validate hippocampal knockout of MR using the in Emx1 Cre: MR fl/fl mice. Scale bar = 50 μm. **b**. Quality control measures in spots from the hippocampal cluster. Distribution of hippocampal read counts per spot, number of genes per spot, and percentage of mitochondrial DNA per spot are shown, with each color representing an animal (3 WT and 3 KO).

**Extended Data Fig. 2.**
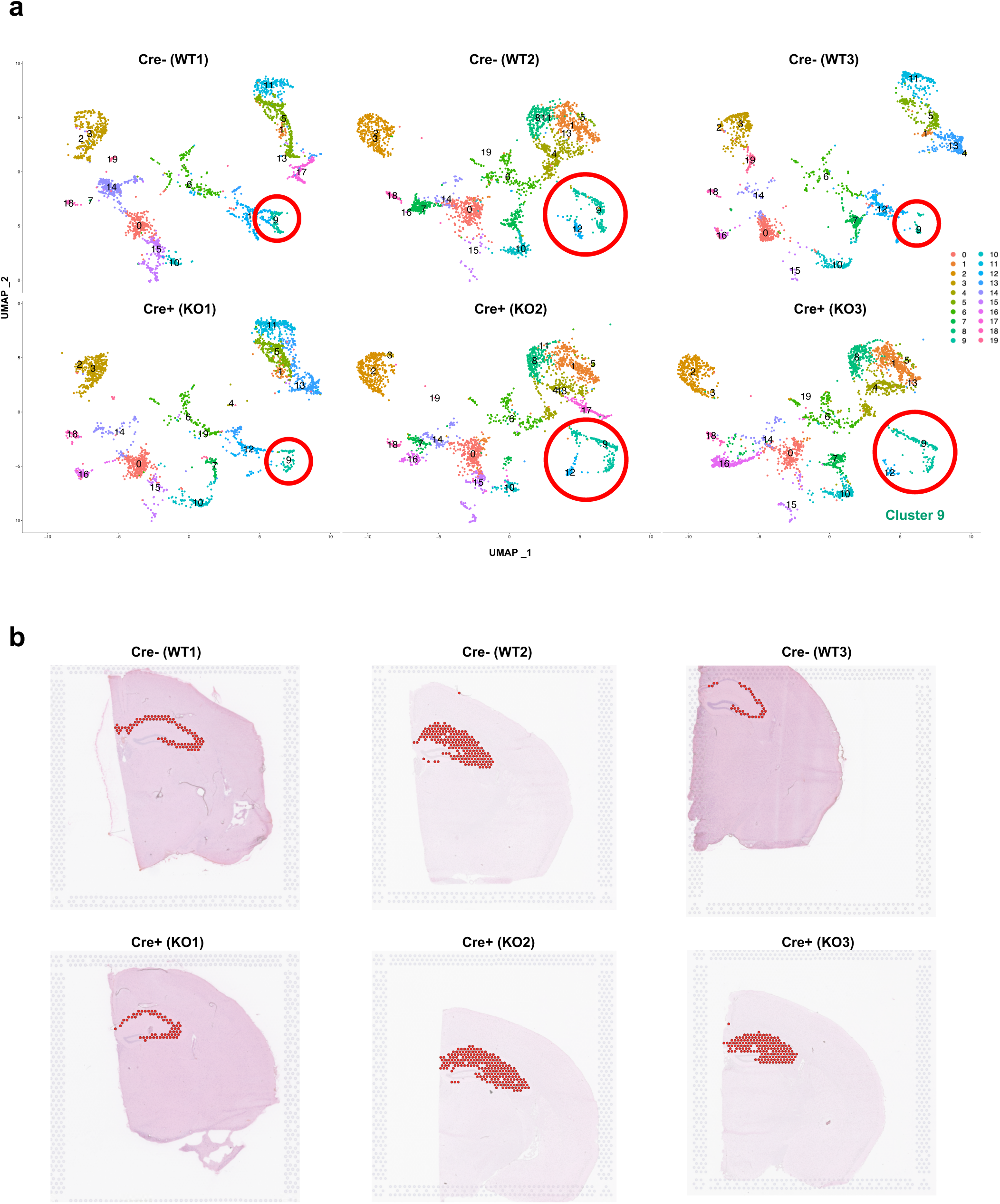
**a**. UMAP projection of the brain tissue based on gene expression clusters. A total of 19 clusters were formed based on gene expression profiles in cell populations within each animal. Plots from the 3 WT and 3 MR KO brains are shown. **b**. ‘Cluster 9’, representing the hippocampus, is shown in the 6 brains. Note images of brains in WT3, KO2, and KO3 were flipped horizontally for illustrative purposes. Batch effects, which appeared to include or not include the stratum radiatum, were not strong enough to be removed by correction and likely reflects the strong similarity between the dendritic fields and the corresponding cell bodies^13^.

**Extended Data Fig. 3.**
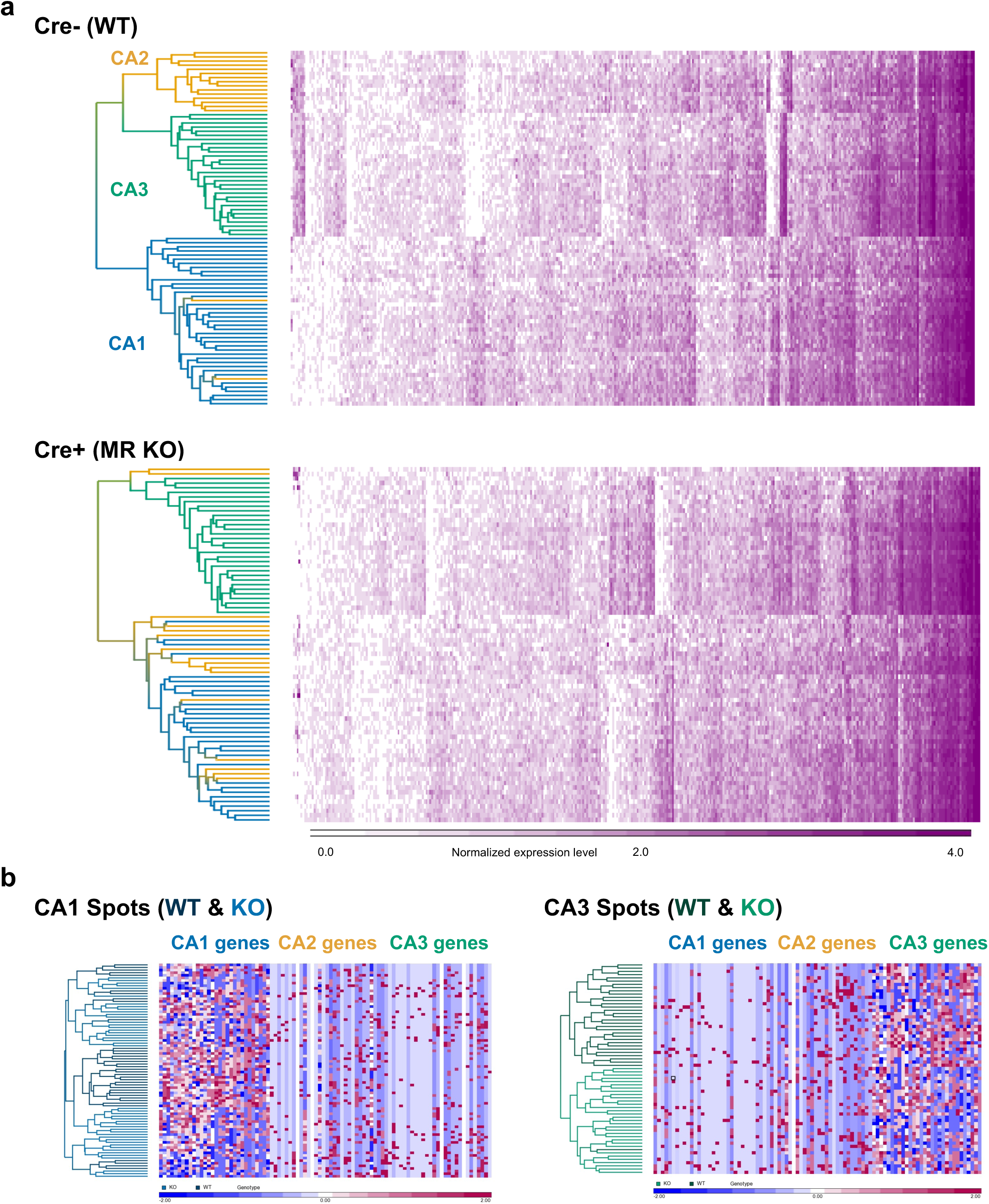
**a**. Dendrograms indicating the hierarchical clustering of spots with similar expression profiles based on 368 hippocampal genes and color-coded according to the spot location. Expression levels are shown as a gradient from white to purple, where expression level zero is white, and high expression is represented by dark purple. “Hippocampal genes” are defined as overexpressed genes in the hippocampal area spots in comparison with the spots in the rest of the areas using a cutoff of FC > 1.2 and adjusted p value < 0.05. **b**. Heatmap of gene expression values for CA1 and CA3 spots from WT and MR KO males for the top 30 CA1 genes, CA2 genes, and CA3 genes as identified by differential expression analysis. Blue = lower expression, Red = higher expression

**Extended Data Fig. 4.**
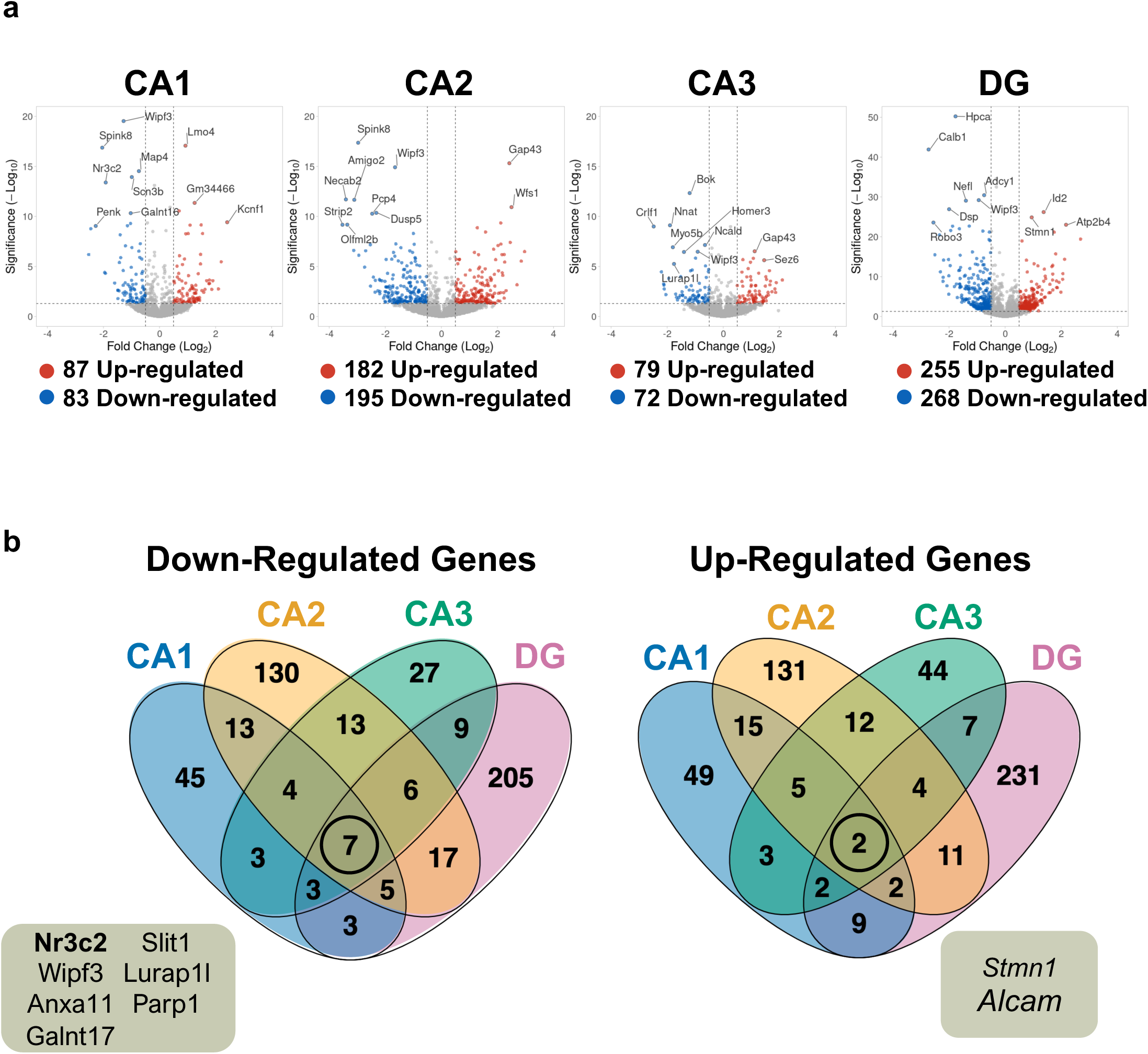
**a**. Volcano plots showing differentially expressed genes between Cre-WT and Cre+ MR KO in CA1, CA2, CA3, and DG. The top 10 genes such as *Nr3c2, Wfs1, Crlf1* and *Calb1* are plotted with Manhattan distance Significance (–log10) = 1.3 (FDR p<0.05) and fold change (Log2) greater than 1 (red) or less than -1 (blue). **b**. Venn diagrams demonstrating minimal overlap of DEGs between hippocampal regions. We found 130 down-regulated genes and 131 up-regulated genes that were unique to area CA2. DEGs common to all 4 regions are listed below each diagram. Notably, *Nr3c2* is down-regulated in all regions.

**Extended Data Fig. 5.**
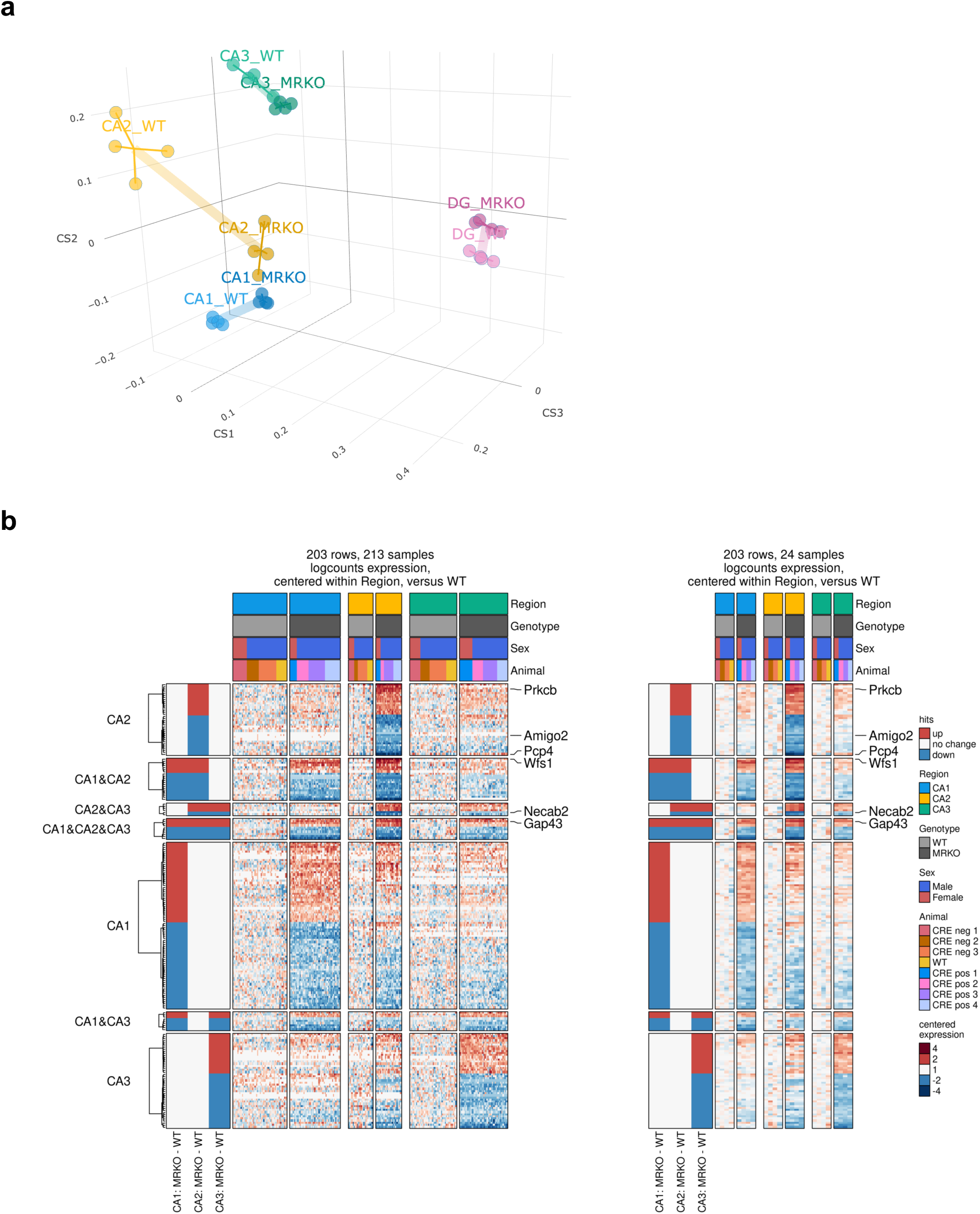
a. Principal component analysis (PCA) highlighting the transcriptional changes associated with MR knockout (KO) shows a shift in the transcriptional profile of CA2 in Cre+ MR KO mice towards CA1, compared to Cre-mice. In MR KO, the correlation between CA1 and CA2 is enhanced, while the correlation between CA3 and CA2 remains unchanged. n = 8 (6 males, 2 female). **b.** Heatmaps depicting the expression levels of MR KO and wildtype (WT) mice across the CA1, CA2, and CA3 hippocampal regions. Log-transformed counts were normalized both by spot (left) and by animal (right). Statistical significance for comparisons between regions is indicated (P < 0.05, n = 8). Relative expression was calculated by log-transforming the raw counts, followed by centering the data to the mean expression of wildtype samples within each region. This normalization approach isolates the differential expression of MR KO relative to wildtype in each specific hippocampal subregion. n = 8 (6 males, 2 female).

**Extended Data Fig. 6.**
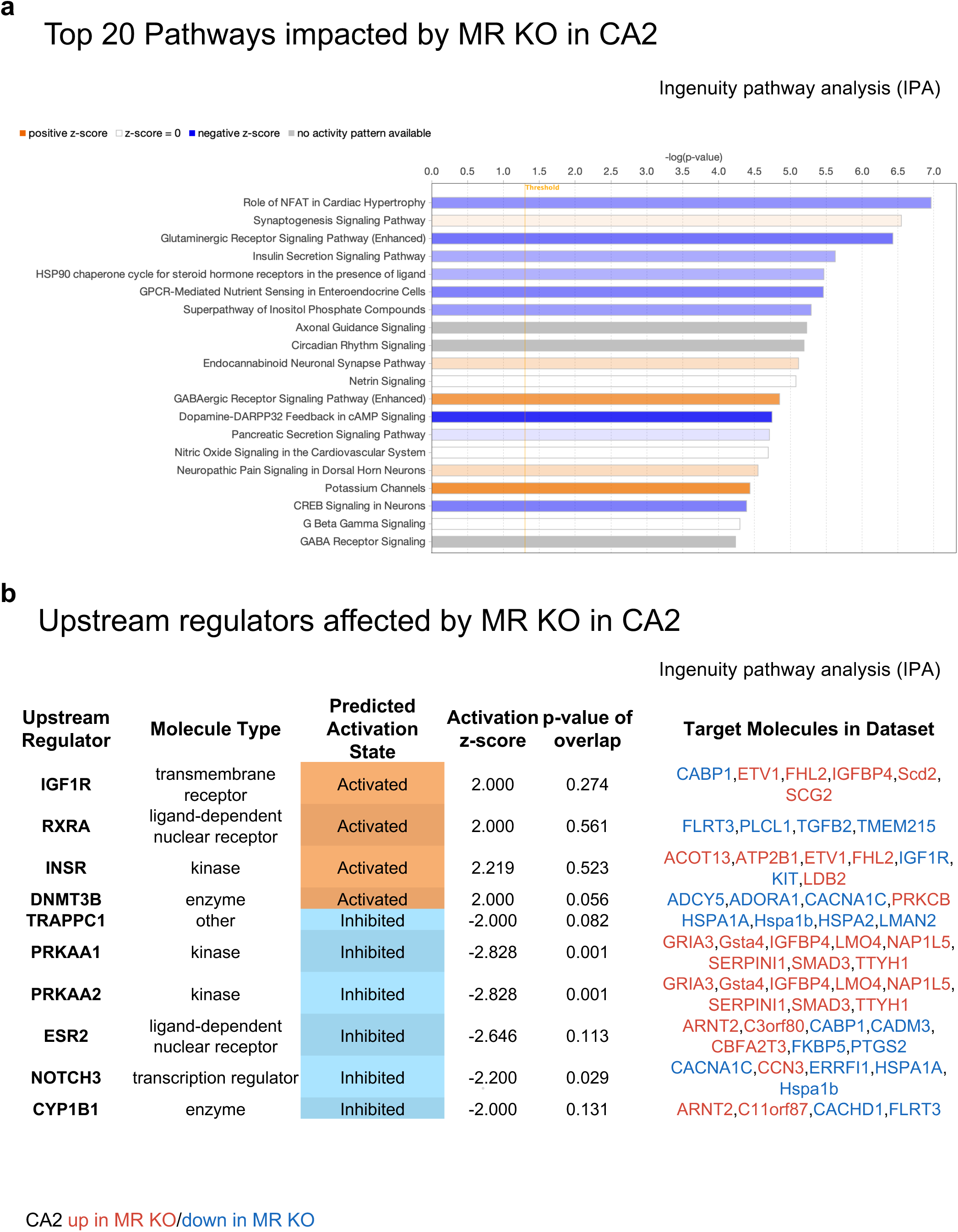
**a**. Differentially expressed genes between Cre-WT and Cre+ MR KO in CA2 with log fold-change values greater than 0.5 or less than -0.5 and FDR <0.05 were included for Ingenuity Pathway Analysis (IPA). Among the top 20 pathways ranked by p-value, the GABAergic receptor signaling and potassium channels pathways are predicted to be activated (orange), while the Dopamine-DARPP32 feedback in cAMP signaling pathway is predicted to be inactivated (blue). The white bars representing netrin, nitric oxide and G Beta Gamma Signaling pathways indicated no clear signal for prediction (z-score = 0). The gray bars representing circadian rhythm, GABA receptor signaling and axon guidance pathways indicated that there is no activity pattern available identified in IPA, despite the highly significant association of the genes within these pathways. **b.** The upstream regulator analysis predicts activation of *IGF1R, RXRA, INSR, and DNMT3B* (z-scores > 2.000), and inactivation of *DNMT3B, TRAPPC1, PRKAA1, PRKAA2, ESR2, NOTCH3,* and *CYP1B1* (z-scores < - 2). Although activation of these regulators is suggested, the high p-value (0.274) for the overlap indicates that this prediction may not be statistically significant. In contrast, the inactivation of *PRKAA1, PRKAA2,* and *NOTCH*3 is supported by statistically significant p-values (< 0.05).

**Extended Data Fig. 7.**
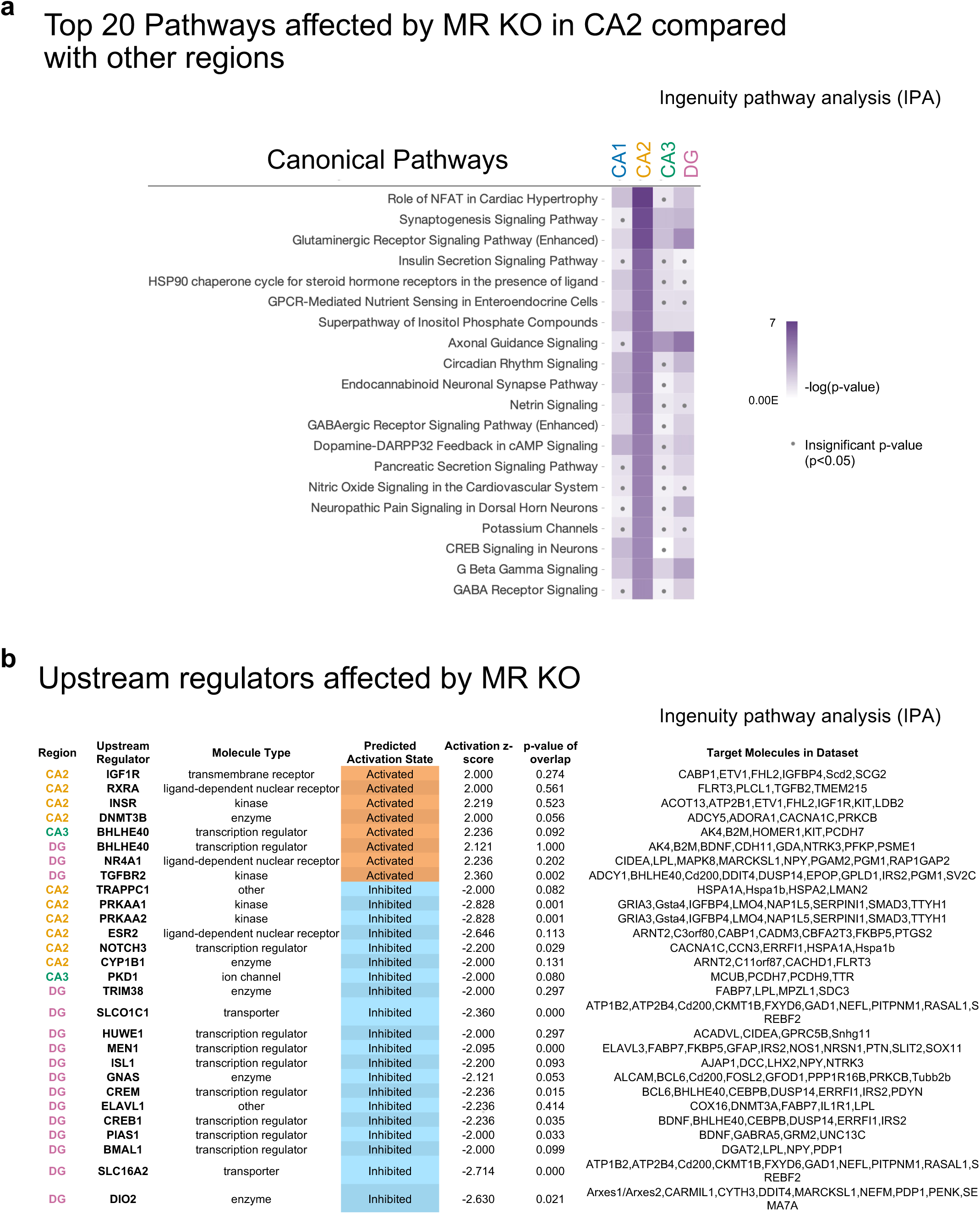
a. Fifteen of the top 20 pathways affected by MR KO in CA2 are not significant in CA3, but are significant in CA1 and DG. Pathways such as glutamatergic receptor signaling, G beta gamma signaling, and the superpathway of inositol phosphate compounds are significant in all regions, with the superpathway of inositol phosphate compounds showing statistically significant inhibition specifically in CA1 and DG. **b.** In the comparison analysis of upstream regulators, no activators or inhibitors were predicted for the CA1 region. In the CA3 region, *BHLEH40* is predicted as an inhibitor and PKD1 as an activator, both with insignificant predictions. In the DG region, several inhibitors, including *SLCO1C1, MEN1, CREM, CREB1, PIAS1, SLC16A2,* and *DIO2*, are statistically significantly predicted, while *TGFBR2*, predicted as an activator, is also statistically significant.

**Extended Data Fig. 8.**
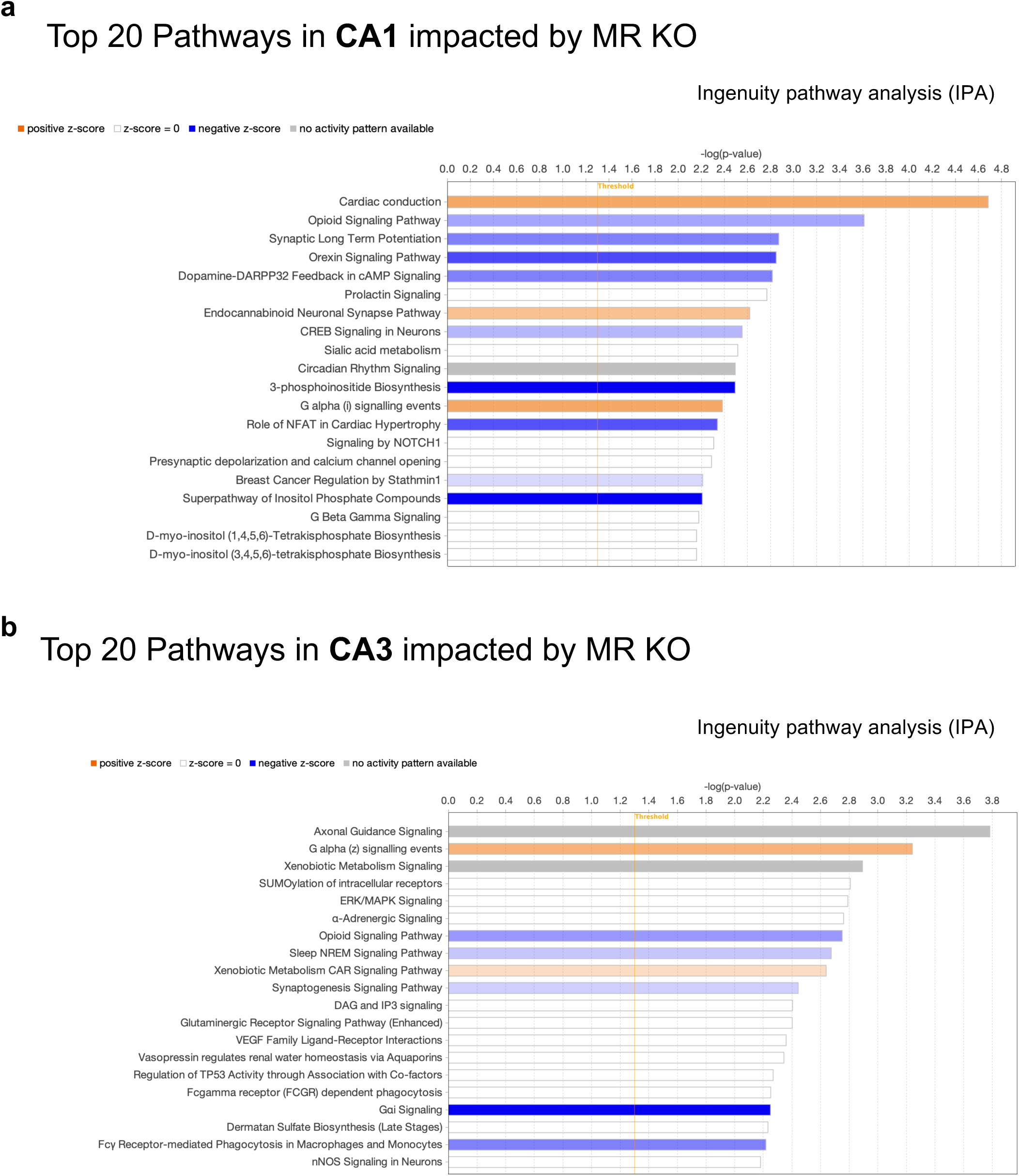
Differentially expressed genes between Cre-WT and Cre+ MR KO in CA1 **(a)** or CA3 **(b)** with log fold-change values greater than 0.5 or less than -0.5 and FDR <0.05 were included for Ingenuity Pathway Analysis (IPA). Among the top 20 pathways in CA1 ranked by p-value, the Cardiac conduction and G alpha (i) signaling event pathways are predicted to be activated (orange), while the 3-phosphoinositide biosynthesis pathway is predicted to be inactivated (blue). The white bars indicated no clear signal for prediction (z-score = 0). The gray bars representing circadian rhythm, xenobiotic metabolism signaling and axon guidance pathways indicated that there is no activity pattern available identified in IPA, despite the highly significant association of the genes within these pathways.

